# KaKs_Calculator 3.0: calculating selective pressure on coding and non-coding sequences

**DOI:** 10.1101/2021.11.25.469998

**Authors:** Zhang Zhang

## Abstract

KaKs_Calculator 3.0 is an updated toolkit that is capable for calculating selective pressure on both coding and non-coding sequences. Similar to the nonsynonymous/synonymous substitution rate ratio for coding sequences, selection on non-coding sequences can be quantified as non-coding nucleotide substitution rate normalized by synonymous substitution rate of adjacent coding sequences. As testified on empirical data, it shows effectiveness to detect the strength and mode of selection operated on molecular sequences, accordingly demonstrating its great potential to achieve genome-wide scan of natural selection on diverse sequences and identification of potentially functional elements at whole genome scale. The package of KaKs_Calculator 3.0 is freely available for academic use only at https://ngdc.cncb.ac.cn/biocode/tools/BT000001.

## Introduction

Detecting natural selection on molecular sequences is of fundamental significance in molecular evolution, comparative genomics and phylogenetic reconstruction, which can provide profound insights for revealing evolutionary processes of molecular sequences and unveiling complex molecular mechanisms of genome evolution [1]. In principle, estimating selection on DNA sequences requires a reference set of substitutions that is free from selection. As synonymous substitutions do not provoke amino acid changes due to the degeneracy of the genetic code, they are expected to be invisible to selection and thus widely used as a reference that reflects the neutral rate of evolution [2]. Consequently, the ratio of nonsynonymous substitution rate (Ka or d_N_) to synonymous substitution rate (Ks or d_S_), namely, ω=Ka/Ks (or d_N_/d_S_), is widely adopted to differentiate neutral mutation (ω≈1) from negative (purifying) selection (ω<1) and positive (adaptive) selection (ω>1), accordingly providing a powerful tool for illuminating molecular evolution of coding sequences (see a popular package in [3]).

Nowadays, a growing body of evidence have shown that non-coding sequences, historically thought as “junk” due to few knowledge on their function relative to coding sequences, are recognized as functional elements to play important regulation roles in multiple biological processes [4] and associate closely with various human diseases [5-7]. Albeit less conserved by comparison with coding sequences, a larger number of non-coding sequences have been identified highly conserved across mammalian genomes [8-10]. Importantly, more non-coding sequences are subject to positive selection and negative selection than previously believed, and particularly, long non-coding RNA (lncRNA) sequences do experience natural selection [11]. As a result, several methods have been proposed for computational detection of selection acting on non-coding sequences [12], which primarily differ in how to choose a reference of unconstrained evolution, such as, synonymous substitutions of neighboring coding gene [13], intron sequences [14, 15] and ancestral repeats [16]. However, there lacks of an implemented algorithm to detect the strength and mode of selective pressure on non-coding sequences, particularly considering an increasing number of non-coding studies conducted worldwide. More importantly, an integrated toolkit that is capable to detect selection on both coding and non-coding sequences is highly desirable, which would help users achieve genome-wide scan of natural selection on diverse sequences.

Towards this end, here we present KaKs_Calculator 3.0, an updated toolkit for calculating selective pressure on both coding and non-coding sequences. Compared with previous versions [17, 18], we implement an algorithm in KaKs_Calculator 3.0 that employs synonymous sites of adjacent coding sequences as a reference to estimate selective pressure acting on non-coding sequences. We test it on empirical data and demonstrate its utility in diagnosing the strength and form of molecular evolution.

### Algorithm

The major update of KaKs_Calculator 3.0 is to incorporate an algorithm that is capable for estimating selective pressure on non-coding sequences. Specifically, it uses synonymous substitutions as a reference baseline (similar to [13]), which, albeit thought to be under weak selection [19-21], has been widely adopted for determining the strength and type of selection operated on coding sequences [22-27]. Similarly, selective pressure on non-coding sequences (ξ) can be quantified as non-coding nucleotide substitution rate (Kn) normalized by neutral substitution rate (assumed as Ks), viz., ξ = Kn/Ks, where Ks is inferred from adjacent coding sequences. As the number of observed substitutions is less than the number of real substitutions, we adopt a nucleotide substitution model (e.g., JC/K2P/HKY) to correct multiple substitutions of non-coding sequences. Taking the HKY model [28] as an example, therefore, Kn can be deduced from the observed transitional and transversional substitutions (S and V, respectively) as well as four nucleotide frequencies (π_A_, π_T_, π_G_ and π_C_), according to Equation 1 (see Equations 1.27 and 1.28 in [29]).

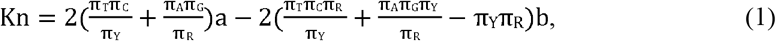

Where 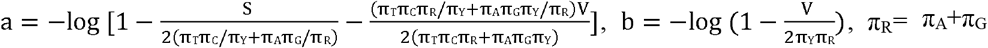 and π_Y_= π_T_+ π_C_ To detect and quantify selection on non-coding sequences, KaKs_Calculator 3.0 provides users with two ways to obtain the value of neutral mutation rate or Ks, which is either calculated from adjacent coding sequences uploaded by users or just specified in a straightforward manner by users (Figure 1). As a consequence, KaKs_Calculator 3.0 is capable to detect selection on both coding and non-coding sequences.

**Figure 1.**
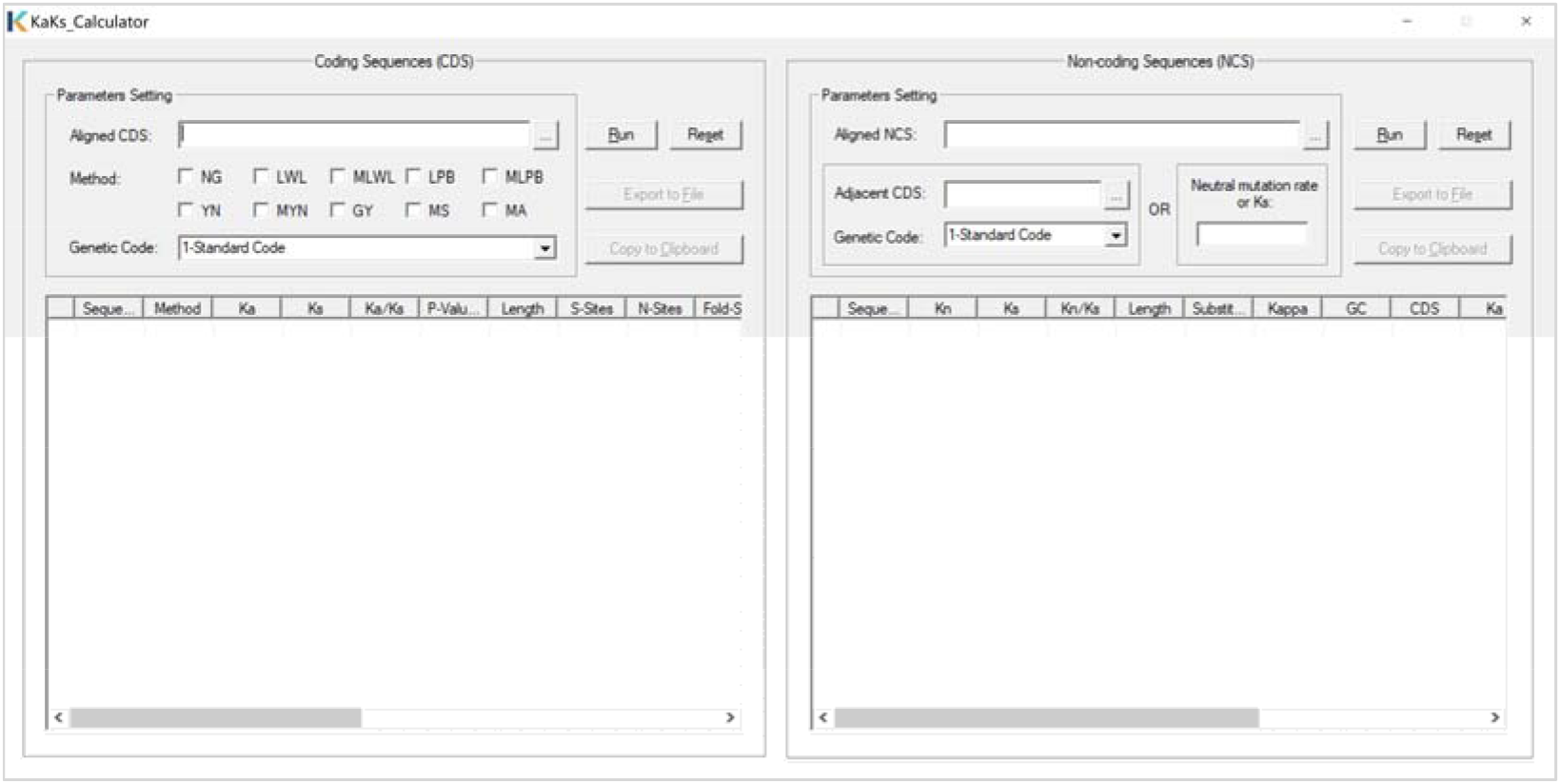
Graphical user interface of KaKs_Calculator 3.0.

KaKs_Calculator 3.0 is implemented in standard C++ language, enabling higher efficiency and easy compilation on different operation systems (Linux/Windows/Mac). In addition to the new functionality for estimating selection on non-coding sequences as mentioned above, it is also updated by fixing bugs and errors. The package of KaKs_Calculator 3.0, including compiled executables, a GUI (graphical user interface) application on Windows, source codes and example data, accompanying with detailed instructions and documentation, is freely available for academic use only at BioCode (https://ngdc.cncb.ac.cn/biocode/tools/BT000001), an open-source platform for archiving bioinformatics tools in the National Genomics Data Center [30], China National Center for Bioinformation.

### Application on empirical data

To test KaKs_Calculator 3.0, we choose three empirical lncRNA genes that are extensively studied according to LncRNAWiki [7] and collect their human-mouse orthologs as well as their adjacent coding orthologs from NGDC LncBook [31] and NCBI RefSeq [32]. Specifically, these non-coding and coding gene symbols with accession numbers are: (1) *H19* (NR_002196.2 vs. NR_130973.1) and *MRPL23* (NM_021134.4 vs. NM_011288.2); (2) *MALAT1* (NR_002819.4 vs. NR_002847.3) and *SCYL1* (NM_020680.4 vs. NM_001361921.1); and (3) *HOTAIR* (NR_003716.3 vs. NR_047528.1) and *HOXC12* (NM_173860.3 vs. NM_010463.2). Based on these orthologous genes, we obtain their corresponding aligned sequences by MAFFT [33] (using parameters: --maxiterate 1000 --localpair). It is noted that one non-coding sequence may have multiple adjacent coding genes, which are completely specified by users and thus can lead to different estimates of Ks and ξ.

By contrasting non-coding nucleotide substitution rate to adjacent synonymous substitution rate, we reveal that, although the coding genes undergo strong purifying selection (ω<1), these three non-coding genes present diverse selective pressure. Strikingly, *HOTAIR* (*Hox* transcript antisense intergenic RNA) exhibits positive selection (ξ>1), whereas the rest two genes experience negative selection (ξ<1). *HOTAIR* is a ∼2.3 kb intergenic RNA transcribed from the antisense strand of the *HOXC* gene cluster [34]. The result of positive selection detected on *HOTAIR* relative to *HOXC12* is consistent well with previous findings that *HOTAIR* evolves faster than the neighbouring genes [35]. On the contrary, *MALAT1* (metastasis-associated lung adenocarcinoma transcript 1), a ∼8.7 kb non-coding RNA flanked by the highly conserved kinase-like gene *SCYL1*, is ubiquitously expressed in almost all human tissues, evolutionarily conserved across mammalian species [36] and associated with various cancers [37]. Thus, ξ=0.464 indicates strong selective constraint on *MALAT1*, in accordance with its physiologic and pathophysiological function [38] and conserved RNA structure [39] as documented by previous studies. Likewise, *H19*, a ∼2.3 kb imprinted maternally expressed transcript located near *MRPL23*, is known for close association with Beckwith-Wiedemann Syndrome and also involved in tumorigenesis [40]. Our result shows that *H19* presents stronger selection constraint as indicated by ξ=0.296, conforming well with its conserved sequence and structure [41]. Taken together, KaKs_Calculator 3.0 is effective in estimating natural selection for non-coding sequences, which has the potential to reveal evolutionarily selective pressures operated on diverse molecular sequences.

## Conclusion

KaKs_Calculator 3.0 is significantly updated by achieving the detection of natural selection on non-coding sequences as well as coding sequences. As testified on empirical data, it is of great utility in calculating natural selection on molecular sequences and thus identifying potentially functional elements at genome-wide scale.

## Code availability

KaKs_Calculator 3.0 is freely available for academic use only at https://ngdc.cncb.ac.cn/biocode/tools/BT000001.

## CRediT author statement

**Zhang Zhang:** Conceptualization, Methodology, Software Development, Supervision, Writing,Reviewing and Editing

## Acknowledgements

I would like to extend special thanks to Lina Ma for constructive suggestions and discussions on this work and Zhao Li for valuable help on data collection. I also thank Zhuojing Fan for designing the logo as well as Qing Guo and Lin Dai for fixing a bug on Windows GUI. I am extremely grateful to a number of users for reporting bugs and sending comments since the first release of KaKs_Calculator in 2006. This work was supported by the Strategic Priority Research Program of the Chinese Academy of Sciences [XDA19050302], the National Natural Science Foundation of China [32030021; 31871328], National Key R&D Program of China [2017YFC0907502], and International Partnership Program of the Chinese Academy of Sciences [153F11KYSB20160008].

---

**Table 1.**
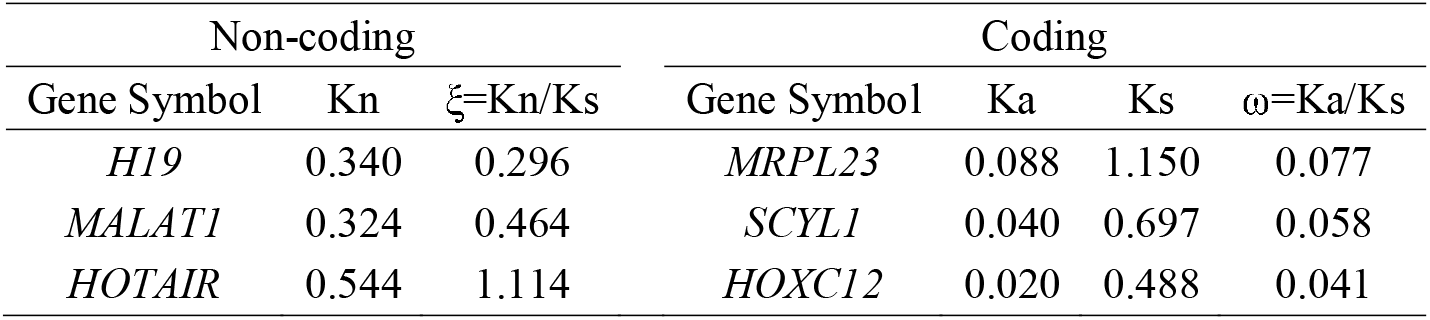
Estimates of selective pressure as well as substitution rates in human-mouse orthologs

| Non-coding |  |  | Coding |  |  |  |
| --- | --- | --- | --- | --- | --- | --- |
| Gene Symbol | Kn | $\xi=K_n/K_s$ | Gene Symbol | Ka | Ks | $\omega=K_a/K_s$ |
| *H19* | 0.340 | 0.296 | *MRPL23* | 0.088 | 1.150 | 0.077 |
| *MALAT1* | 0.324 | 0.464 | *SCYL1* | 0.040 | 0.697 | 0.058 |
| *HOTAIR* | 0.544 | 1.114 | *HOXC12* | 0.020 | 0.488 | 0.041 |

## References

[1] Li W-H. Molecular Evolution. Sunderland, Massachusetts: Sinauer Associates, 1997.

[2] Hurst LD. The Ka/Ks ratio: diagnosing the form of sequence evolution. Trends Genet 2002;18:486.

[3] Yang Z. PAML 4: phylogenetic analysis by maximum likelihood. Mol Biol Evol 2007;24:1586–91.

[4] Dunham I, Kundaje A, Aldred SF, Collins PJ, Davis CA, Doyle F, et al. An integrated encyclopedia of DNA elements in the human genome. Nature 2012;489:57–74.

[5] Luo H, Bu D, Shao L, Li Y, Sun L, Wang C, et al. Single-cell Long Non-coding RNA Landscape of T Cells in Human Cancer Immunity. Genomics Proteomics Bioinformatics 2021.

[6] Anastasiadou E, Jacob LS, Slack FJ. Non-coding RNA networks in cancer. Nat Rev Cancer 2018;18:5–18.

[7] Liu L, Li Z, Liu C, Zou D, Li Q, Feng C, et al. LncRNAWiki 2.0: a knowledgebase of human long non-coding RNAs with enhanced curation model and database system. Nucleic Acids Res 2022;doi: 10.1093/nar/gkab998.

[8] Bejerano G, Pheasant M, Makunin I, Stephen S, Kent WJ, Mattick JS, et al. Ultraconserved elements in the human genome. Science 2004;304:1321–5.

[9] Habic A, Mattick JS, Calin GA, Krese R, Konc J, Kunej T. Genetic Variations of Ultraconserved Elements in the Human Genome. OMICS 2019;23:549–59.

[10] Guttman M, Amit I, Garber M, French C, Lin MF, Feldser D, et al. Chromatin signature reveals over a thousand highly conserved large non-coding RNAs in mammals. Nature 2009;458:223–7.

[11] Diederichs S. The four dimensions of noncoding RNA conservation. Trends Genet 2014;30:121–3.

[12] Zhen Y, Andolfatto P. Methods to detect selection on noncoding DNA. Methods Mol Biol 2012;856:141–59.

[13] Wong WS, Nielsen R. Detecting selection in noncoding regions of nucleotide sequences. Genetics 2004;167:949–58.

[14] Parsch J, Novozhilov S, Saminadin-Peter SS, Wong KM, Andolfatto P. On the utility of short intron sequences as a reference for the detection of positive and negative selection in Drosophila. Mol Biol Evol 2010;27:1226–34.

[15] Hoffman MM, Birney E. Estimating the neutral rate of nucleotide substitution using introns. Mol Biol Evol 2007;24:522–31.

[16] Bush EC, Lahn BT. A genome-wide screen for noncoding elements important in primate evolution. BMC Evol Biol 2008;8:17.

[17] Zhang Z, Li J, Zhao XQ, Wang J, Wong GK, Yu J. KaKs_Calculator: calculating Ka and Ks through model selection and model averaging. Genomics Proteomics Bioinformatics 2006;4:259–63.

[18] Wang D, Zhang Y, Zhang Z, Zhu J, Yu J. KaKs_Calculator 2.0: a toolkit incorporating gamma-series methods and sliding window strategies. Genomics Proteomics Bioinformatics 2010;8:77–80.

[19] Shabalina SA, Spiridonov NA, Kashina A. Sounds of silence: synonymous nucleotides as a key to biological regulation and complexity. Nucleic Acids Res 2013;41:2073–94.

[20] Hershberg R, Petrov DA. Selection on codon bias. Annu Rev Genet 2008;42:287–99.

[21] Plotkin JB, Kudla G. Synonymous but not the same: the causes and consequences of codon bias. Nature Reviews Genetics 2011;12:32–42.

[22] Li WH, Wu CI, Luo CC. A new method for estimating synonymous and nonsynonymous rates of nucleotide substitution considering the relative likelihood of nucleotide and codon changes. Mol. Biol. Evol. 1985;2:150–74.

[23] Nei M, Gojobori T. Simple methods for estimating the numbers of synonymous and nonsynonymous nucleotide substitutions. Mol. Biol. Evol. 1986;3:418–26.

[24] Li WH. Unbiased estimation of the Rates of synonymous and nonsynonymous substitution. J. Mol. Evol. 1993;36:96–9.

[25] Goldman N, Yang Z. A codon-based model of nucleotide substitution for protein-coding DNA sequences. Mol. Biol. Evol. 1994;11:725–36.

[26] Yang Z, Nielsen R. Estimating Synonymous and Nonsynonymous Substitution Rates Under Realistic Evolutionary Models. Mol. Biol. Evol. 2000;17:32–43.

[27] Zhang Z, Li J, Yu J. Computing Ka and Ks with a consideration of unequal transitional substitutions. BMC Evol Biol 2006;6:44.

[28] Hasegawa M, Kishino H, Yano T. Dating of the human-ape splitting by a molecular clock of mitochondrial DNA. J Mol Evol 1985;22:160–74.

[29] Yang Z. Computational Molecular Evolution. London: Oxford University Press, 2006.

[30] CNCB-NGDC Members & Partners. Database Resources of the National Genomics Data Center, China National Center for Bioinformation in 2021. Nucleic Acids Res 2021;49:D18–D28.

[31] Ma L, Cao J, Liu L, Du Q, Li Z, Zou D, et al. LncBook: a curated knowledgebase of human long non-coding RNAs. Nucleic Acids Res 2019;47:D128–D34.

[32] Li W, O’Neill KR, Haft DH, DiCuccio M, Chetvernin V, Badretdin A, et al. RefSeq: expanding the Prokaryotic Genome Annotation Pipeline reach with protein family model curation. Nucleic Acids Res 2021;49:D1020–D8.

[33] Katoh K, Standley DM. MAFFT multiple sequence alignment software version 7: improvements in performance and usability. Mol Biol Evol 2013;30:772–80.

[34] Tang Q, Hann SS. HOTAIR: An Oncogenic Long Non-Coding RNA in Human Cancer. Cell Physiol Biochem 2018;47:893–913.

[35] He S, Liu S, Zhu H. The sequence, structure and evolutionary features of HOTAIR in mammals. BMC Evol Biol 2011;11:102.

[36] Gutschner T, Hammerle M, Diederichs S. MALAT1 -- a paradigm for long noncoding RNA function in cancer. J Mol Med (Berl) 2013;91:791–801.

[37] Meseure D, Vacher S, Lallemand F, Alsibai KD, Hatem R, Chemlali W, et al. Prognostic value of a newly identified MALAT1 alternatively spliced transcript in breast cancer. Br J Cancer 2016;114:1395–404.

[38] Zhang X, Hamblin MH, Yin KJ. The long noncoding RNA Malat1: Its physiological and pathophysiological functions. RNA Biol 2017;14:1705–14.

[39] Smith MA, Gesell T, Stadler PF, Mattick JS. Widespread purifying selection on RNA structure in mammals. Nucleic Acids Res 2013;41:8220–36.

[40] Hurst LD, Smith NG. Molecular evolutionary evidence that H19 mRNA is functional. Trends Genet 1999;15:134–5.

[41] Juan V, Crain C, Wilson C. Evidence for evolutionarily conserved secondary structure in the H19 tumor suppressor RNA. Nucleic Acids Res 2000;28:1221–7.

